# Genetic and chemical differentiation of *Campylobacter coli* and *Campylobacter jejuni* lipooligosaccharide pathways

**DOI:** 10.1101/050328

**Authors:** Alejandra Culebro, Joana Revez, Ben Pascoe, Yasmin Friedmann, Matthew D Hitchings, Jacek Stupak, Samuel K Sheppard, Jianjun Li, Mirko Rossi

**Affiliations:** Department of Food Hygiene and Environmental Health, Faculty of Veterinary Medicine, University of Helsinki, Agnes Sjöbergin katu 2, Helsinki, FI-00014, Finland.; College of Medicine, Institute of Life Science, Swansea University, Singleton Park, Swansea SA2 8PP, United Kingdom.; Institute for Biological Sciences, National Research Council, Ottawa, Ontario K1A 0R6, Canada.; Department of Biology and Biochemistry, University of Bath, Bath, United Kingdom

**Keywords:** Campylobacter, LOS, glycoconjugate, Qui3*p*NAcyl

## Abstract

Despite the importance of lipooligosaccharides (LOS) in the pathogenicity of campylobacteriosis, little is known about the genetic and phenotypic diversity of LOS in *C. coli*. In this study, we investigated the distribution of LOS locus classes among a large collection of unrelated *C. coli* isolates sampled from several different host species. Furthermore, we paired *C. coli* genomic information and LOS chemical composition for the first time to identify mechanisms consistent with the generation of LOS phenotypic heterogeneity. After classifying three new LOS locus classes, only 85% of the 144 isolates tested were assigned to a class, suggesting higher genetic diversity than previously thought. This genetic diversity is at the basis of a completely unexplored LOS structure heterogeneity. Mass spectrometry analysis of the LOS of nine isolates, representing four different LOS classes, identified two features distinguishing *C. coli* LOS from *C. jejuni*’s. GlcN-GlcN disaccharides were present in the lipid A backbone in contrast to the GlcN3N-GlcN backbone observed in *C. jejuni*. Moreover, despite that many of the genes putatively involved in Qui3*p*NAcyl were absence in the genomes of various isolates, this rare sugar was found in the outer core of all *C. coli*. Therefore, regardless the high genetic diversity of LOS biosynthesis locus in *C. coli*, we identified species-specific phenotypic features of *C. coli* LOS which might explain differences between *C. jejuni* and *C. coli* in terms of population dynamics and host adaptation.

**Depositories (where applicable):** The whole genome sequences of *C. coli* are publicly available on the RAST server (http://rast.nmpdr.org) with guest account (login and password ‘guest’) under IDs: 195.91, 195.96-195.119, 195.124-195.126, 195.128-195.130, 195.133, 195.134, 6666666.94320

## INTRODUCTION

Campylobacteriosis is the most common bacterial food-borne disease in developed countries, with over 200,000 human cases reported annually in the European Union alone (1). The true burden of the disease in the population is likely underestimated, as many infections result in mild gastroenteritis (1). Approximately ~80% of reported infections are caused by *Campylobacter jejuni* and 7-18% of cases are attributed to *C. coli*. Therefore *C. coli*, is among the five most important bacterial aetiological agents of human gastroenteritis (2, 3).

As in other Gram-negative bacteria, *Campylobacter* spp. cell surface glycoconjugates, including lipooligosaccharides (LOS), play an important role in serum and bile resistance, resistance to phagocytic killing, adhesion, invasion, and survival in host cells (4-8). Current knowledge on LOS diversity has been based primarily on work in *C. jejuni* and its role in promoting severe clinical symptoms (9-12). *C. jejuni* LOS is a potent TLR4 agonist and the subsequent immune response is affected by changes in LOS structure and composition (10-14). Additionally, due to molecular mimicry between human gangliosides and certain LOS structures, *C. jejuni* has been identified as one of the causative agents of the Guillain-Barré syndrome (15). Valuable insights into the genetic origins of significant strain variable traits have been gained by studying the effect that *C. jejuni* LOS genotypes have on phenotype (16-21). However, little is known about the variability of LOS in *C. coli* despite of the impact of this structure on human health. Only two studies have addressed the variation in gene composition in the *C. coli* LOS biosynthesis locus, describing nine genetic classes composed of a variable combination of 10 to 20 genes (22, 23). Three further studies have explored the chemical composition of LOS in *C. coli* and a single structure has been described (24-27).

We investigated the diversity and distribution of LOS locus classes among a large collection of unrelated *C. coli* isolates sampled from several different host species. By analysing genomic data with the LOS chemical composition of selected isolates, we identified mechanisms that may be responsible for the observed difference in LOS phenotype. We expanded the current LOS typing scheme by including three additional LOS locus classes. Despite the extensive introgression between *C. coli* and *C. jejuni* (28, 29), only negligible levels of recombination were detected in LOS biosynthesis genes, which might explain the distinctive species-specific chemical features observed herein.

## METHODS

### Bacterial isolates, cultivation, and DNA extraction

In total, 144 *C. coli* isolates, including 90 isolated from swine, 34 from humans, 18 from poultry, and two from wild birds, were chosen for LOS locus screening. The selection comprised 133 *C. coli* isolates from previous studies collected between 1996 and 2012 from Finnish human patients, chicken and pigs reared in Finland, and wild birds sampled in Helsinki region (22, 30-36). These were augmented by 11 *C. coli* isolates from the Campynet (CNET) collection (hosted by DSMZ GmbH, https://www.dsmz.de/). Isolate selection was based on genotype (PFGE, MLST), host, country of origin, and year of isolation to encompass the largest possible diversity. Cultivation and DNA isolation were carried out as previously described (22), unless otherwise stated.

### PCR

The length of LOS biosynthesis loci were determined by amplifying the region between orthologue 10 (LOS biosynthesis glycosyltransferase, *waaV*) and orthologue 16 (one or two domains glycosyltransferases) (ID numbers according to Richards and colleagues (23)). PCR reactions were set up as follows: 25 μl reactions containing 0.5 U Phusion high-fidelity (Thermo Scientific), 200 μM of each dNTP (Thermo Scientific), 0.4 μM of each primer (ORF3F2 and waaV; Table 1), 1 X Phusion GC buffer (Thermo Scientific), 700 μM of MgCl2 (Thermo Scientific), and 5 μl of template. Cycling conditions were as follows: one cycle at 98 °C for 30 s followed by 30 cycles of denaturation at 98 °C for 10 s, annealing at 62.4 °C for 30 s, extension at 72 °C for 6 min, and a final elongation at 72 °C for 6 minutes. The size of the LOS locus was estimated by gel electrophoresis with 1 kb-plus (Thermo Scientific) and long-range (Thermo Scientific) molecular weight markers. Specific primers for each class, based on the previously described *C. coli* LOS locus classes (I to VIII) (23). Typing scheme is summarised in Table 1. Since global alignment using progressiveMauve (37) revealed that LOS locus class IV and V (23) differ by only 3 single nucleotide polymorphism (which resulted in fragmentation of orthologue 1959 in class V), hereafter the two LOS locus classes are considered as a single class named IV/V. Isolates with unexpected LOS size, negative to all tested orthologues, or with unexpected combination of orthologues profile, were classified as untypable.

**Table 1.**
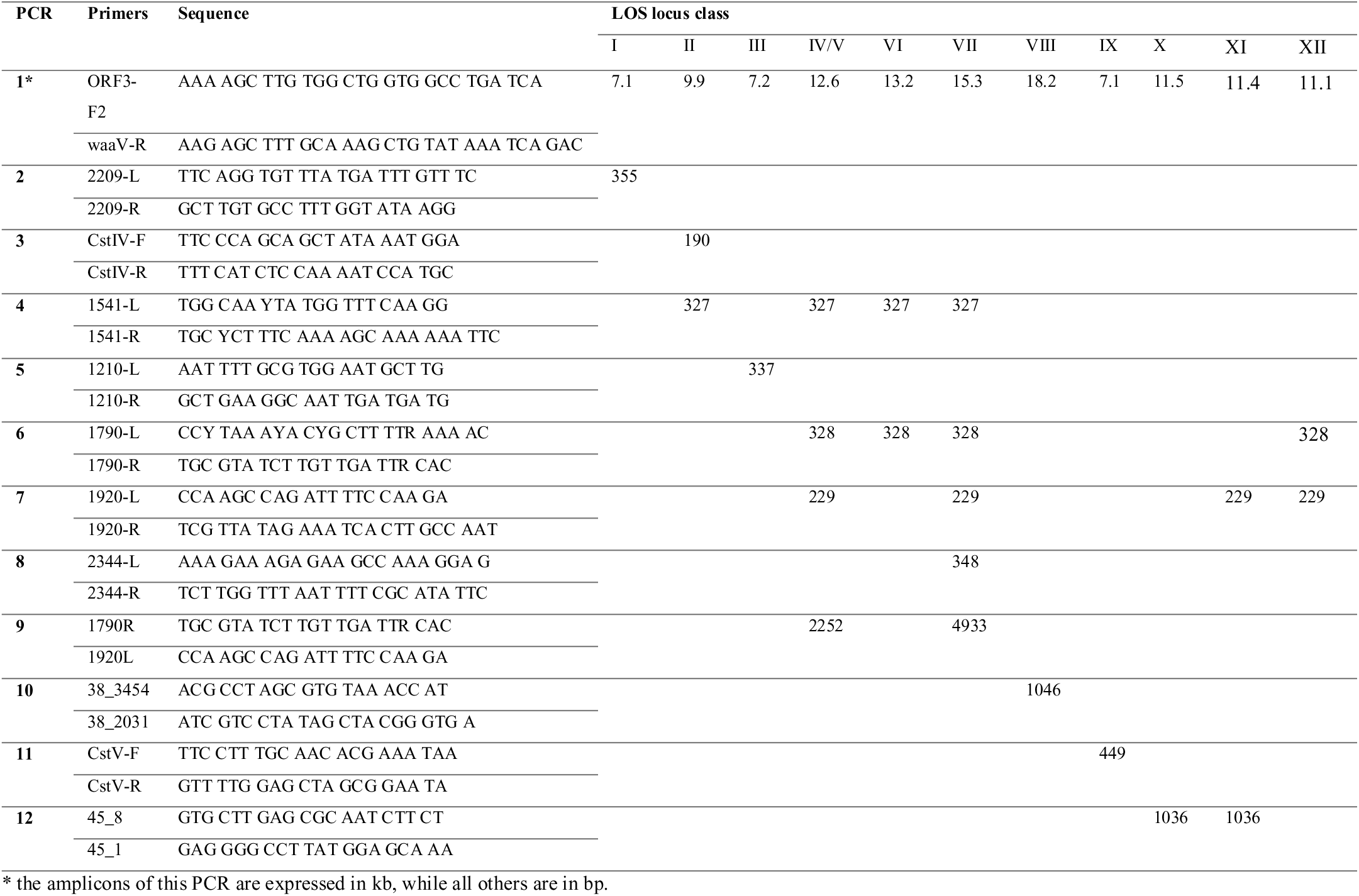
List of primers used in the present study and expected sizes of the amplicons.

### Genome sequencing and annotation

For ascertaining the LOS locus classes, 35 isolates were chosen for genome sequencing using either HiSeq or MiSeq. For HiSeq, NGS library preparation, enrichment, sequencing, and sequence analyses were performed by the Institute for Molecular Medicine Finland (FIMM Technology Centre, University of Helsinki, Finland). MiSeq sequencing was performed by Institute of Life Science, Swansea University (Swansea, United Kingdom). Reads were filtered and assembled using SPAdes Assembler v. 3.3.0 (38). Primary annotation of all the genomes was performed using Rapid Annotation using Subsystem Technology (RAST) (39). Sequences were manually curated using Artemis (40) and LOS locus classes were aligned and compared with ACT (41).

### Orthologue clustering and phylogenetic analysis

A database including all the translated coding sequences of *C. jejuni* and *C. coli* LOS biosynthesis was assembled (23) using Richards and colleagues (23) orthologues nomenclature. Reciprocal all-versus-all BLASTp search was performed (threshold E ≤ 1e-10) (42) and orthologous groups were determined by orthAgogue and MCL (ignoring E-values, percent match length ≥ 80% and inflation value of 5 (43, 44)). The groups of orthologues (GOs) were then aligned using MUSCLE and back-translated to nucleotide sequence using Translatorx perl script (45-47). Maximum likelihood phylogenetic reconstruction of each GO was performed in MEGA6.06 (48) using Kimura-2 as nucleotide substitution model and a discrete Gamma distribution (4 categories) to model evolutionary rate differences among sites. A total of 100 bootstrap runs were performed which were summarized in a 95% consensus tree.

### LOS silver staining

LOS profiles were assessed by silver staining as described earlier (49), with some modifications. In brief, the absorbance of the biomass obtained from a 16 h Nutrient broth n°2 (Oxoid) culture (100 rpm, microaerobic atmosphere, 37 °C) was adjusted to an OD600 of 0.5. Cells were digested with 20 mg/ml proteinase K (Thermo Scientific), and incubated at 55 °C for 1 h followed by boiling for 10 min. Samples were then diluted 1: 5 in loading buffer, and ran in 15% SDS gels. Gels were silver stained for visualization.

### CE-MS and EA-OTLC-MS analyses

Biomass was produced in broth as indicated above and LOS was prepared with the rapid method applying microwave irradiation as described previously (50). In short, the lyophilized biomass was suspended in 50 μl of 20 mM ammonium acetate buffer (pH 7.5) containing DNase (100 μg/ml) and RNase (200 μg/ml) and heated by direct microwave irradiation. Proteinase K was then added to a final concentration of 60 μg/ml and heated under the same conditions. Solutions were allowed to cool at room temperature and subsequently dried using a Speed Vac (vacuum centrifuge concentrator; Savant). LOS samples were washed three times with methanol (100 μl) with vigorous stirring. Insoluble residues were collected by centrifugation and resuspended in 30 μl water for electrophoresis-assisted open-tubular liquid chromatography-electrospray MS (EA-OTLC-MS) analysis. A sheath solution (isopropanol-methanol, 2:1) was delivered at a flow rate of 1.0 μL/minute. Separation was performed using 30 mM morpholine in deionized water, pH 9.0. A separation voltage of 20 kV, together with a pressure of 500 mbar, was applied for the EA-OTLC-MS analysis. The electrospray ionization (ESI) voltage applied on the sprayer was set at −5.2 kV. Data acquisition was performed for an m/z range of 600 to 2000 at a 2s/spectrum scan rate.

### Statistical analysis

Fisher’s exact test was used to assess the association of LOS locus class with host. P values below 0.05 were considered significant.

## RESULTS

### PCR typing method for *C. coli* LOS locus diversity

We explored the genetic diversity of LOS biosynthesis in 144 *C. coli* isolates (Supplementary table 1) using a PCR typing scheme based on published LOS locus class definitions (2, 3). Isolates were classified into putative LOS locus classes according to their PCR-profile and LOS locus size as described in Table 1. The LOS PCR typing scheme was validated by genome sequencing of 35 isolates. Typing results are summarised in Table 2. We were able to designate 68% of the isolates to one of the nine previously published LOS locus classes (22, 23). Most of the isolates were assigned to LOS locus class II (17%) with the remaining isolates assigned to LOS classes IV/V (15%), III (13%), VI (13%), VIII (7%), I (2%), VII (1%), and IX (0.7%). The final 46 (out of 144, ~32%) isolates remained untypable by this method (Table 1).

**Table 2.** Distribution of LOS classes among hosts

| LOS class | Total (%) | Human | Swine | Poultry | Wild birds |
| --- | --- | --- | --- | --- | --- |
| I | 3 | 2 | 0 | 1 | 0 |
| II | 24 (17) | 7 | 13 | 4 | 0 |
| III | 18 (13) | 4 | 13 | 0 | 1 |
| IV/V | 22 (15) | 3 | 16 | 3 | 0 |
| VI | 18 (13) | 1 | 15 | 2 | 0 |
| VII | 2 (1) | 1 | 1 | 0 | 0 |
| VIII | 10 (7) | 7 | 1 | 2 | 0 |
| IX | 1 | 1 | 0 | 0 | 0 |
| X | 22 (15) | 3 | 18 | 1 | 0 |
| XI | 1 | 0 | 1 | 0 | 0 |
| XII | 1 | 0 | 0 | 0 | 1 |
| Untypable | 22 (15) | 5 | 12 | 5 | 0 |

Six untypable isolates, with a LOS locus length of ~11.5 kbp, were sequenced (45, 63, 114, 125, 149, and 153). All isolates belong to the novel LOS locus class X. This new class shares 12 (out of 15) orthologues with other LOS locus classes (see below), and is characterised by the presence of three unique genes. A search of the NCBI database with amino acid sequences, revealed sequence similarity with: (i) hypothetical protein of *Helicobacter* sp. MIT 05-5293 (e-value 1e^−98^; identity 45%); (ii) hypothetical protein of *Helicobacter hepaticus* (e-value 3e^−108^; identity 53%); (iii) UDP-N-acetylglucosamine 2-epimerase of *H. hepaticus* (e-value 3e^−165^; identity 63%). Following this finding, primers were designed (Table 1) for LOS locus class X which identified a further 15% of the isolates, from all sources (Table 2). The genomes of isolates 138 and 99, which have a similar LOS size to class X but a different PCR profile (Table 1) were also sequenced. Analysis of these genomes found two additional LOS locus classes, defined as XI (isolate 138) and XII (isolate 99). In total, we were able to assign a LOS locus class to 85% of the isolates in our collection by incorporating these additional classes. LOS profile diversity was high, suggesting that further LOS locus classes may be described in the future.

### Origin of the novel LOS locus classes X, XI, and XII

As in *C. jejuni, C. coli* exhibits a mosaic LOS loci (19) with several classes containing similar orthologous loci. LOS locus classes X and XI are very similar to each other, diverging only at a single locus. These classes also have similarity in gene content and organisation to LOS locus classes I, III, IV/ V, V I, an d VII (Fig. 1). The genetic context of LOS locus class X in orthologues 16, 1850, and 1668 is unclear, while loci organisation of isolates 45, 63, and 114 is similar to class III, isolates 125 and 149 show similarity to LOS locus class I (Fig. 1). Extensive recombination and gene reorganisation between classes may be masking the origin of common shared loci (Fig. 1). Orthologue 1821 in LOS locus class X may originate from LOS locus class VI, while the LOS locus class XI allele is more closely related to that of class IV/V. Orthologue 1967 in isolates 45, 125, 149, and 153 originated from the same common ancestor, while in isolates 63 and 114 appears to originate from LOS locus class VI. LOS locus class XII shares genes with LOS locus classes I, IV/V, VII and IX, and it is characterized by the presence of a set of unique genes having the best BLASTp hit against NCBI nr with: (i) methyltransferase type 12 of *H. hepaticus* (e-value 6e^−75^; identity 58%); (ii) hypothetical protein of *Anaerovibrio lipolyticus* (e-value 5e^−102^; identity 65%); (iii) phosphoserine phosphatase of *Helicobacter* sp. MIT 05-5293 (e-value 3e^−92^; identity 63%) (Fig. 1). Proposed functions for each ORF of the herein newly identified LOS classes are described in Supplemental Table 2.

**Figure 1.**
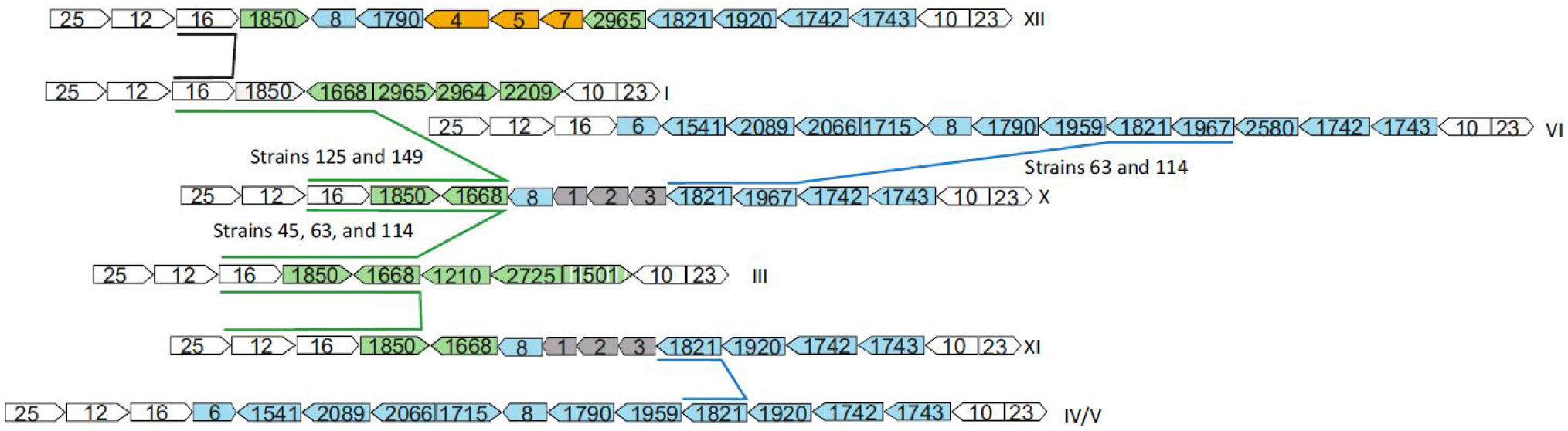
LOS locus classes related to X, XI, and XII. Arrows represent ORFs. Genes coloured white are common to all LOS classes. Genes coloured green are present in class I and/or III. Genes coloured blue are present in classes IV/V and VI. Grey genes are common among classes X and XI. The orange genes are particular of the class XII. Striped genes are fragmented. Gene size is not drawn to scale.

### Cluster analysis of the LOS locus classes

Both species share a total of 19 LOS orthologues (23) and with previous evidence of introgression between *C. coli* and *C. jejuni* in mind (28, 29) we attempted to quantify the level of interspecies recombination in *C. coli* LOS diversity. We compared individual gene descriptions of the LOS loci rather than the original gene family ontologies used by Richards and colleagues (23). Out of the 19 shared orthologues, 16 gene locus descriptions split into species-specific clusters while only three were common in both species (orthologues 10, 16 and 1821). Interspecies gene transfer was investigated by comparing the topology of individual gene trees with the overall population structure (22). Evidence of interspecies gene transfer was only observed for gene 10 (23) (lipooligosaccharide biosynthesis glycosyltransferase, *waaV*) where all *C. coli* loci of LOS locus class II formed a monophyletic clade with *C. jejuni* genes (Fig. 2). Thus, interspecies recombination likely only has a limited effect on the LOS loci diversity observed in *C. coli*.

**Figure 2.**
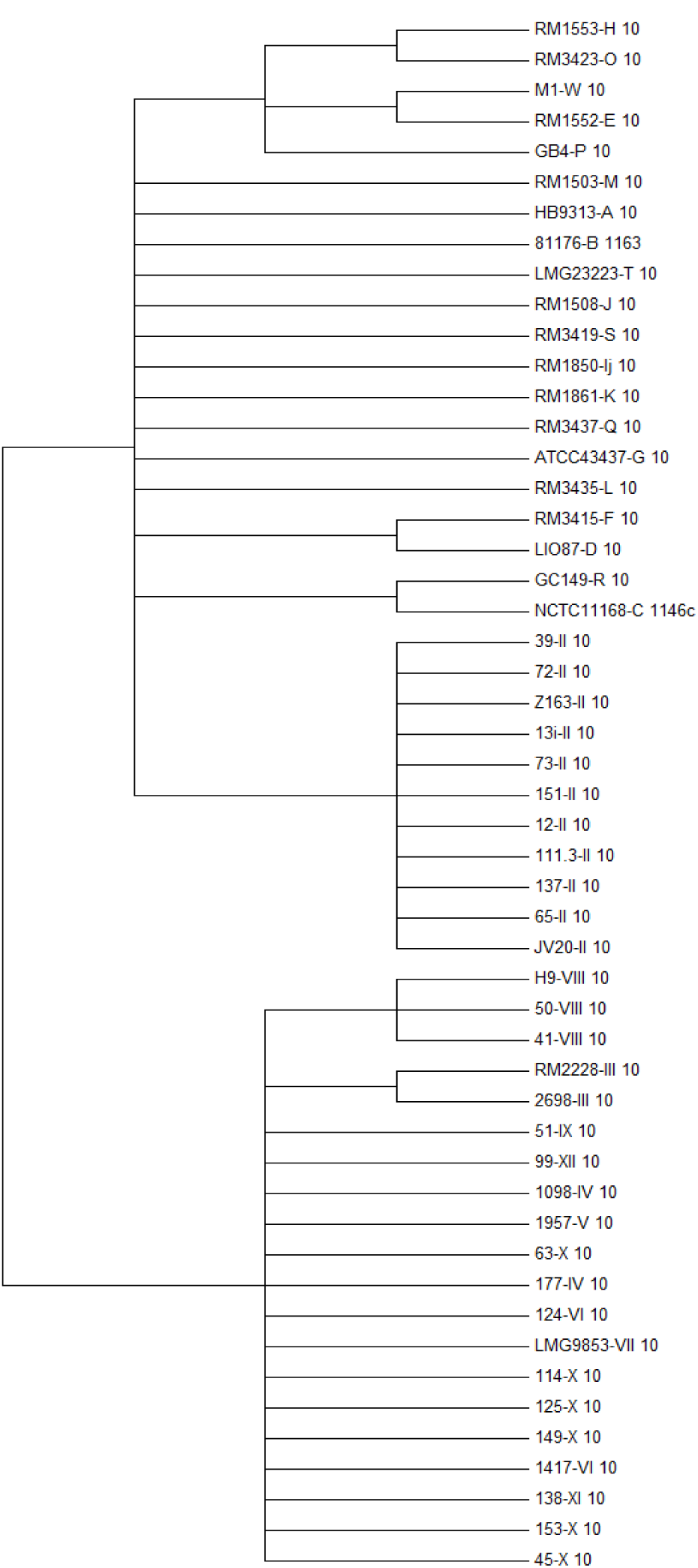
Consensus cladogram representing the evolutionary relationship among orthologues belonging to GO 10 (nomenclature from Richard *et al*. 23). The 95% bootstrap consensus tree was built from 100 replicates.

### Host-LOS locus class association

The distribution of LOS locus classes was not random between hosts and was investigated further, supplemented with data from Richards and colleagues (23). The distribution of LOS locus classes by source of isolation is represented in Figure 3. All LOS locus classes, except in class XII, were present among strains isolated from humans. More than half (57%) of the clinical isolates were LOS locus classes II, III and VIII, while LOS locus classes VI, VII and X were found less often in clinical cases. Most pig isolates were of LOS locus class X, but also frequently found among LOS locus classes II, III, IV/V, and VI. Only one pig isolate belonged to LOS locus class VIII and no pig strain was from classes I, IX, or XII. Poultry isolates were also found among all LOS locus classes, except for classes VII, IX, and XII. Most poultry isolates were classified as LOS locus class II.

**Figure 3.**
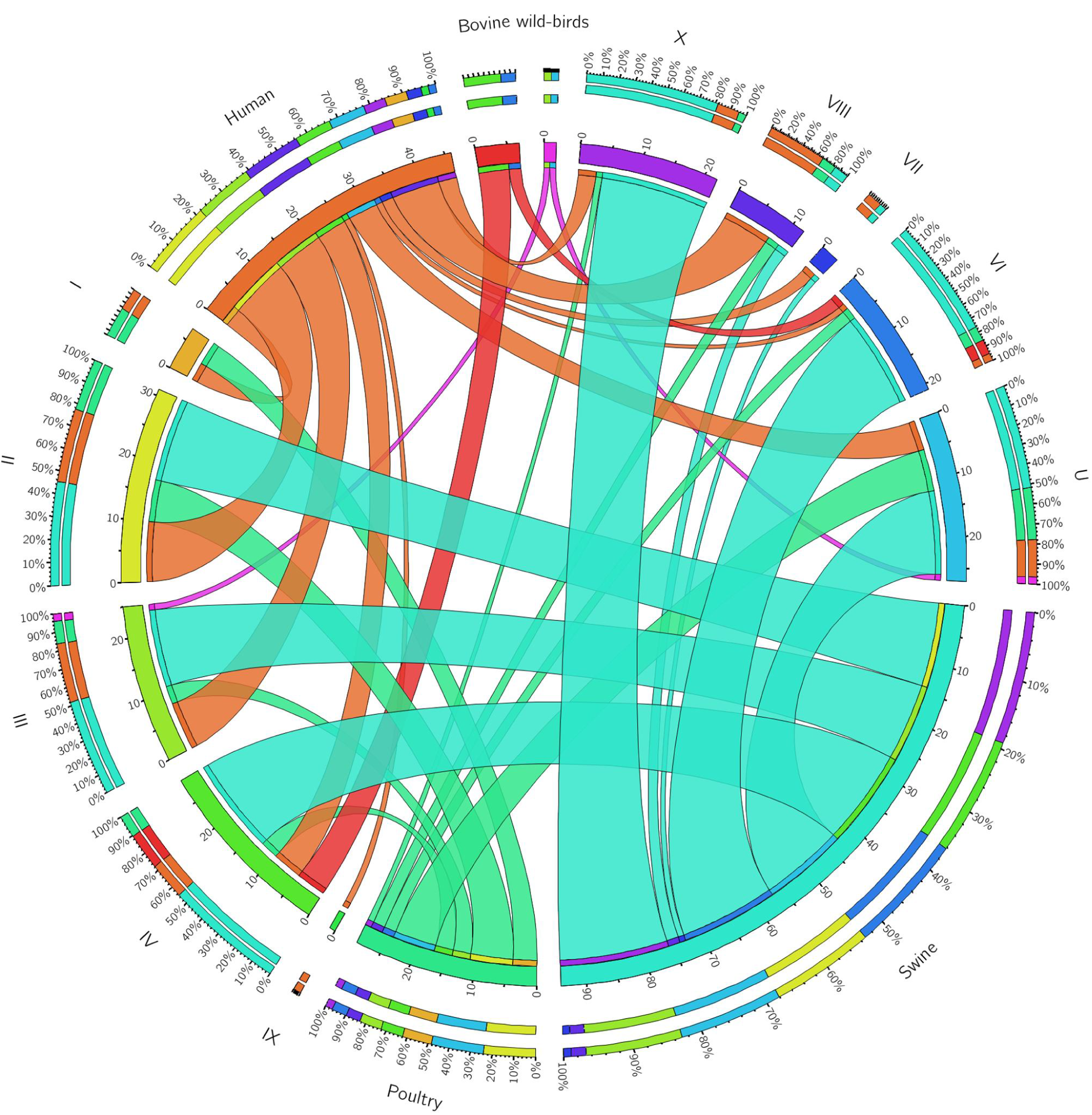
Visualization of the distribution of LOS locus classes of *C. coli* strains isolated from different hosts, from both our collection and those from Richards and colleagues (23).

There was a positive association (p <0.05) of class VIII to human clinical infection, while class VI was negatively associated with clinical infection. Swine was positively associated to classes VI and X, but negatively associated to classes I and VIII. Poultry was positively associated only to LOS locus class I. Bovine and wild-bird isolates were underrepresented in the dataset; however, some association was observed in bovine (class IV/V) and wild bird isolates (class XII). Isolates carrying LOS locus classes II and III were equally distributed among humans, pigs, and poultry.

### Chemical analysis of *C. coli* LOS composition

Silver staining SDS PAGE gels of LOS extracts provided migration profiles for nine isolates from classes II, VIII, X, and XI (Fig. 4A). A complimentary mass spectroscopy approach was used (CE-MS and EA-OTLC-MS) (Examples of MS for strains 65 and 137 are shown on Supplemental Fig. 1) to explore inter-and intra-LOS class structural diversity. The oligosaccharide (OS) composition of each of the nine isolates was predicted based on the fragment ions and components of the previously reported *C. coli* OS (24). Size and composition of the lipid A group was defined for each glycoform by tandem mass spectrometry. *C. coli* isolates exhibited a hexa-acylated lipid A containing four tetradecanoic (14:0) and two hexadecanoic (16:0) acid chains, modified with two phosphate residues (51-53). Only 2-amino-2-deoxy-d-glucose (GlcN) disaccharides were detected in *C. coli* isolates, in contrast to the hybrid backbone of β-1’-6 linked 3-diamino-2, 3-dideoxy-D-glucopyranose (GlcN3N) and GlcN observed in *C. jejuni* (51, 53). This results in a lipid A with two esters and two amide-linked acyl chains.

**Figure 4.**
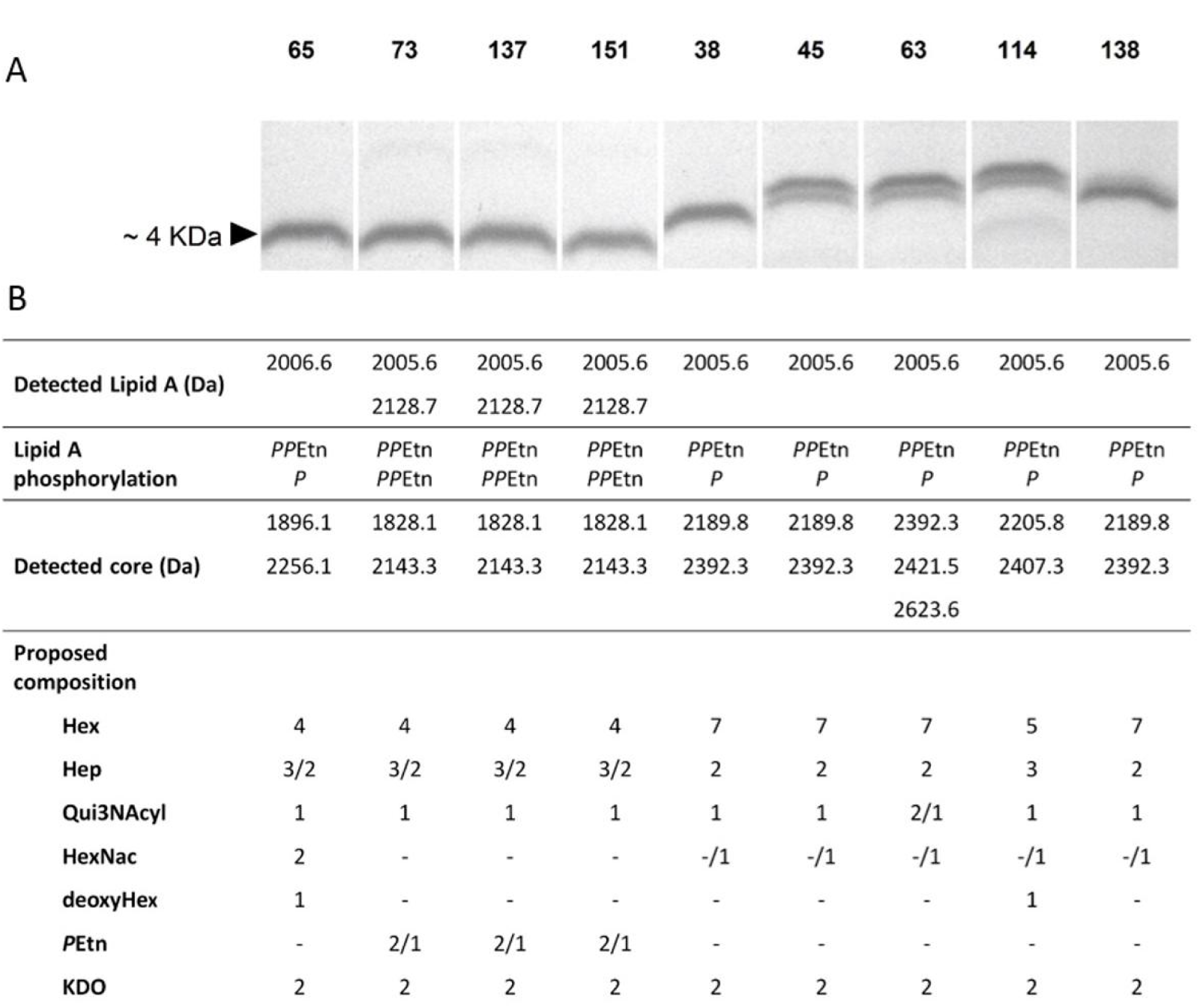
*C.coli* LOS biochemical profiles. **A**) Silver-stained LOS. **B**) Proposed chemical composition based on MS and MS/MS results analysis of intact LOS.

The two detected lipid A forms differed by 123 Da, which corresponds to a second phosphoethanolamine (PEtn) residue. The lowest mass (2005 Da) lipid A was detected in all samples, while LOS class II isolates (except for isolate 65) had an additional lipid A species with 2 PEtn residues (2128.7 Da). As in *C. jejuni, C. coli* exhibited a conserved inner core consisting of two L-glycero-D-manno-heptose (Hep) residues attached to a 3-deoxy-D-manno-octulosonic residue (Kdo) which is linked to the lipid A through a Kdo linker (17, 53). In the variable outer core region at least one Qui*p*3NAcyl residue (where Qui*p*3NAc represents 3-acylamino-3,6-dideoxy-D-glucose in which the N-acyl residue was a 3-hydroxybutanoyl) was detected in all isolates. Although more than one OS was detected by MS in all isolates (Fig. 4B), only isolates from LOS locus classes X and XI exhibited visible high-M_r_ and low - M _r_ LOS on SDS-PAGE (Fig. 4A).

Intra-LOS class diversity was observed in both LOS class II and class X. Isolate 65 displayed a LOS composition like other LOS class II isolates but with the addition of two hexosamines (HexNac) and one deoxyhexose (deoxyHex), and absence of PEtn residues (Fig. 4B). Likewise, isolates 45 and 63 shared similar LOS composition, with the exception of a variable Qui*p*3NAcyl residue in isolate 63. In contrast, isolate 114 exhibited a very different LOS composition compared with other isolates of the same class, including the presence of a third Hep and a deoxyHex as well as a reduced number of hexoses (Hex) (Fig. 4B).

### Genetic and phenotypic diversity within *C. coli* LOS class II

The four strains with LOS locus class II shared 99.64% DNA sequence similarity and from 99.39% to 99.98% pairwise alignment identity. Isolate 65 was the most dissimilar among strains with LOS locus class II due to deletion of large fragments. Deletions resulted in shorter 2400 and 2473 orthologues, as one pseudogene (Fig. 5). Orthologues 2470 and 2471 were also truncated as one pseudogene (re-annotated as 2470-1), as evidenced by isolate 151. The remainder of the class II isolates had an insertion of 68 nt in 2470-1, disrupting the orthologue (Fig. 5). Despite the differences observed in orthologue 2470-1 isolates 73, 137, and 151 had identical LOS chemical composition.

**Figure 5.**
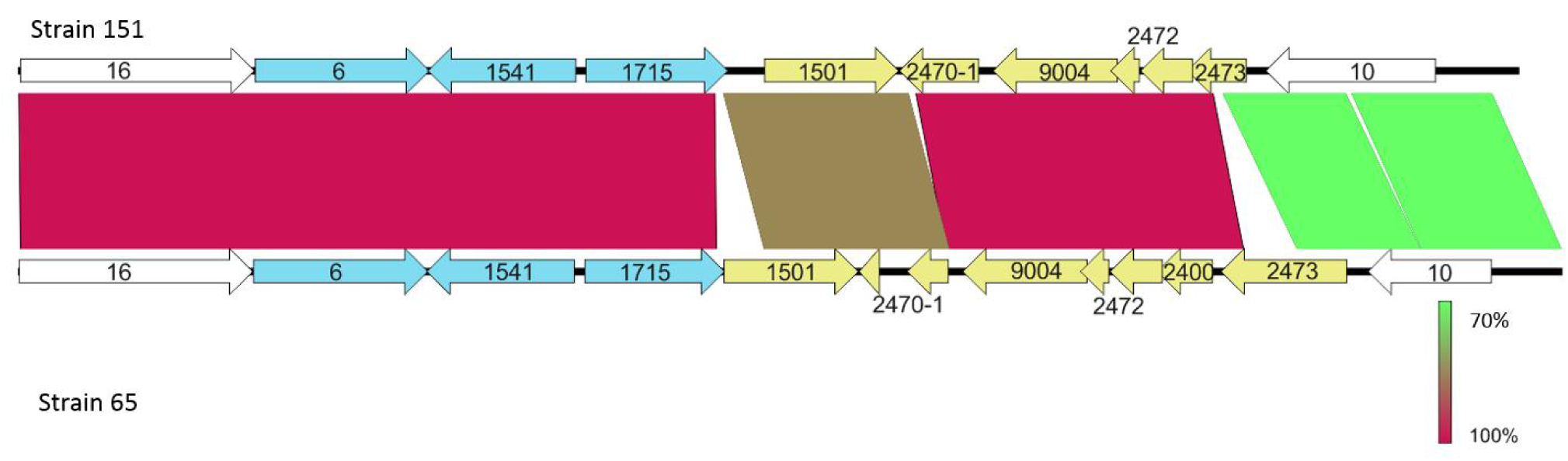
Comparison of nucleotide sequence of LOS locus class II strains 151 and 65. Genes coloured white are common to all LOS classes. Genes coloured blue are present in LOS locus classes IV/V, VI, and VII. Yellow coloured genes are particular to LOS locus class II.

Amino acid sequences of orthologues 6, 1541, 1501, 2472, and 10 were identical (100%), while orthologues 9004 and 16 exhibited a single amino acid difference in isolate 65. All isolates exhibited differences in the C-terminal of orthologue 1715 and were variable in the number of Hep and/or PEtn residues observed. However, no GC homopolymeric tracts or other possible genetic signals associated to phase variation were identified within the LOS loci.

### Genetic and phenotypic diversity within *C. coli* LOS locus class X

In LOS locus class X the overall sequence identity among strains was 99.31%, with percentage identity ranging from 98.96% to 99.94% in pairwise alignments, with strain 45 being the most distant. Although some minor gaps were observed, single point mutations were largely responsible for the diversity observed at nucleotide level. The largest insertion (69 nt) was seen in strain 63 between orthologues 2 and 3. Between strains, 100% amino acid identity was observed in orthologues 16, 8, and 2, while one or two amino acid substitutions were present in orthologues 1668, 1, 1821, 1967, and 1743. The most prominent difference was seen in orthologue 1742 in the form of a deleted A base at position 668, resulting in the premature translational termination in isolates 114 and 63. Furthermore, several single amino acid substitutions were detected in orthologue 1742 in strain 45, while 100% identity was observed between isolates 63 and 114. In spite of dissimilar LOS composition, the only difference observed within the LOS locus between isolates 63 and 114 was in eight amino acids at the C-terminal of orthologue 3.

## DISCUSSION

*Campylobacter* LOS is a fundamental feature involved in the pathogenesis of gastroenteritis and post-infection sequelae (10-14, 54, 55). However, despite the burden imposed by *C. coli* and the importance of this structure in campylobacteriosis, little is known about the LOS diversity in this species (23-27). Therefore, we sought to contribute to the paucity of information by investigating the variability and distribution of *C. coli* L OS locus genetic classes in a large collection of isolates and by coupling genomic and LOS chemical composition data for the first time.

We developed a PCR methodology which was able to classify 85% of the isolates into a LOS class (22, 23). Among them, we described three additional LOS locus classes, named X, XI, and XII, which accounted for 17% of the isolates in our collection. The remaining untypable isolates (15%) suggests that further new classes will likely be described in the future and that *C. coli* LOS biosynthesis is more diverse than previously observed (23).

This genetic diversity is at the basis of a completely unexplored LOS structure heterogeneity which might contribute substantially to the population dynamics of *C. coli*, including host specificity. We combined our 144 isolates with 33 *C. coli* previously studied (23) to investigate the non-random distribution of LOS locus classes among different hosts. All hosts were significantly associated with at least one LOS locus class. In particular, isolates possessing LOS locus classes VI and X were predominantly isolated from swine, which have very high prevalence of *C. coli* (up to 99%) (56). Both of these classes were rarely detected in human isolates, which is supported by a previous source attribution study in Scotland in which pigs are a relatively unimportant source of *C. coli* human infections (56). The majority of human cases in our study were assigned to LOS locus classes II or III, which were also found in swine and poultry isolates. However, human isolates we re overrepresented among LOS locus class VIII, which was rarely detected in the sources included in this study. This indicates the presence of other, unknown potential reservoirs contributing to human infections, which is corroborated in a previous study where 54% of human *C. coli* strains were attributed to other sources than poultry and pig (56). In opposition to previous findings (2), we did not observe partitioning between bovine and poultry sourced strains, and LOS locus classes previously shown to be associated with bovine hosts were populated by isolates of poultry and swine origin. Due to the limited number of isolates available from alternative sources, the host-LOS class associations found in this study may not necessarily represent the true *C. coli* population structure in various hosts. However, our findings suggest that generalist isolates possessing LOS locus class II and III might be more successful at colonizing multiple species and, as seen in generalist lineages of *C. jejuni* ST-45 and ST-21 clonal complexes, being largely responsible for human infections (29).

Mosaic *C. coli* LOS classes appear to have arisen by the insertion and/or deletion of genes or gene cassettes through homologous recombination, as previously described in *C. jejuni* (19). Despite of substantial genome-wide introgression between agricultural *C. coli* and *C. jejuni* (22, 28), very limited interspecies recombination was detected among LOS biosynthesis loci. Only orthologue 10 (*waaV*) in *C. coli* LOS locus class II may have originated as result of recombination with *C. jejuni*. These results confirmed previous studies (28), and are corroborated by the species-specific features detected in the chemical composition of *C. coli* LOS.

GlcN disaccharide backbones, which is the most common structure among members of the family *Enterobacteriaceae* (53), were predicted in the lipid A of all analysed *C. coli* strains. This result is in contrast to the hybrid GlcN3N-GlcN backbone observed in *C. jejuni*. The genes *gnnA* and *gnnB*, located outside the LOS biosynthesis locus, are associated to the synthesis of GlcN3N-substituted lipid A (9, 57). Inactivation of either of these genes in *C. jejuni* resulted in the substitution of an *N-*linked with an *O*-linked acyl chain and an increases LOS biological activity in humans (9). *C. coli* contains in a similar genomic location both genes, having approximately 70% BLASTp score ratios against *C. jejuni* orthologues (9). Yet, *C. coli gnnA* and *gnnB* are separated by a putative cobalamin independent methionine synthase II in the same gene orientation. We suggest therefore three possible explanations for the absence of GlcN3N in *C. coli* lipid A backbone: (i) single or multiple mutations in the putative binding sites of *gnnA* and *gnnB* have rendered one or both enzymes inactive, as observed in functional studies in other bacteria (57, 58); (ii) *gnnB-gnnA* operon transcription might be hampered by the presence of the putative methionine synthase II (9); (iii) *gnnA* and *gnnB* may be involved in the biosynthesis of alternative glycoconjugates in *C. coli* (57). Nevertheless, the substitution of an *N-*linked with an *O*-linked acyl chain in *C. coli* might have an impact in host-bacterial interaction and adaptation (9).

A second species-specific feature, common among all our analysed isolates, was the presence of at least one Qui*p*3NAcyl residue. Qui*p*3N is an unusual deoxysugar, which has been observed in the O-antigen of various Gram negative bacteria and in the S-layer of glycoprotein glycans of some Gram positives (59-61). Although rarely studied, Qui*p*3 N has also been found in the OS of LOS class E, H, and P isolates in *C. jejuni* exclusively as an *N-*acetyl derivative (Qui*p*3NAc) (50, 62-64). Conversely, Qui*p*3N has only been reported in *C. coli* as an *N-*acyl derivative with two possible substituents; 3-hydroxybutanoyl or 3-hydroxy-2, 3-dimethyl-5-oxoprolyl (65). The presence of Qui*p*3NAcyl in *C. coli* was first described by Seltmann and Beer (65), and later on it was reported in several *C. coli* (26). However, the molecular basis behind the biosynthesis of this sugar and associated glycoconjugate in *C. coli* remains unknown. The dTDP-D-Qui*p*3NAc biosynthesis pathway has, to our knowledge, only been described in the Gram positive *Thermoanaerobacterium thermosaccharolyticum* (66). This pathway involves five enzymes; a thymydylyltransferase (*rmlA*), a 4, 6-dehydratase (*rmlB*), a 3, 4-isomerase (*qdtA*), a transaminase (*qdtB*), and a transacetylase (*qdtC*). Genome comparison of *T. thermosaccharolyticum* and *C. coli* identified homologs of *rmlA* (GO 1743), *rmlB* (GO 1742), *qdtA* (GOs 1920 and 1967), and *qdtB* (GO 8) in a subset of strains. However, no homologue for *qdtC* was found in *C. coli*. This may be expected as *C. coli* Qui*p*3N is an *N-*acyl derivative instead of an *N-*acetyl derivative found in *T. thermosaccharolyticum* (24, 65). These results are in agreement with previous studies in which *C. jejuni* isolates carrying these orthologues in the LOS locus have been found to express Qui*p*3NAc in their LOS (23, 50, 62-64). Despite the presence of this sugar in all *C. coli* investigated in this study, as described above, the putative dTDP-D-Qui*p*3NAc biosynthesis genes are only present in a subset of strains all belonging to LOS classes IV/V, VI, VII, X, and XI. Furthermore, truncation of orthologue 1742 due to a single base deletion should have resulted in the loss of Qui*p*3NAcyl in isolates 114 and 63, which was not the case. Cross talk between different glycosylation pathways have been previously observed in *C. jejuni* (62, 67). Thus, due to Qui*p*3NAcyl being ubiquitously found in *C. coli* LOS structures, we hypothesize that the synthesis of this residue might be carried out by genes in conserved glycosylation pathways. Because of structural similarity between Qui*p*3NAc and bacillosamine precursors, it is tempting to speculate that the *pgl* system may play a role in the biosynthesis of Qui*p*3NAc in *C. coli*.

In all *C. coli*, phase variation was observed in at least one sugar residue. Phase variation in *C. jejuni* has mainly been associated to repeats of GC homopolymeric tracts (20). However, no GC tracts were detected in the LOS locus of the chemically analysed *C. coli* isolates. Further inspection of all the LOS locus sequences generated in this and previous studies (22, 23) revealed that G-tracts are uncommon in *C. coli* LOS. Only isolates from LOS class IV/V and VI had G-tracts longer than 5 bases in their LOS biosynthesis locus. It is therefore unlikely that the observed phase variation in our analysed samples was due to slipped strand mispairing due to homopolymeric tracts within the LOS locus. These data suggest that other mechanisms, such as post-transcriptional regulation or epigenetic methylation of DNA, might be responsible for phenotypic variation in LOS composition in *C. coli*.

Among LOS locus class II isolates, strain 65 exhibited the most divergent composition. Orthologue 1715 (*wlaTB*) has been associated with a HexNac residue in *C. jejuni* 81116 (62) and the diversity observed in the C-terminal of this orthologue may be responsible for the absence of HexNAc residues in isolates 73, 137, and 151. However, further research is required to confirm the exact role of 1715 in LOS biosynthesis. Similarly to LOS locus class II, strains with LOS locus class X isolates minor genetic dissimilarities resulted in major differences in LOS chemical composition.

Isolates 65 and 114 also contained a deoxyHex residue in the LOS. No orthologues potentially involved in deoxyHex synthesis were identified within the LOS region in isolates 65, suggesting that genes outside the LOS locus may play a bigger role in LOS biosynthesis than previously thought. DeoxyHexoses, such as 6-deoxy-β-l-altrose, fucose or rhamnose have been frequently detected in the O-chain of the lipopolysaccharide (LPS) of several Gram-negative species (68, 69). In the genus *Campylobacter*, these sugars have been described as components of *C. jejuni* capsule (70) and *C. fetus* LPS (71).

In conclusion, the genetic and biochemical diversity of *C. coli* is greater than expected. *C. coli* LOS is characterised by a lipid A consisting of GlcN-GlcN disaccharides and an outer core substituted with at least one Qui*p*3NAcyl residue. Our results hint at cross talk between different glycosylation pathways, which has not been generally considered to play a role in LOS diversity. The relevance of these characteristic features for the ecology and virulence of *C. coli* is yet to be explored.

## Abbreviations

LOS, lipooligosaccharides; RAST, Rapid Annotation using Subsystem Technology; GOs, groups of orthologues; EA-OTLC-MS, electrophoresis-assisted open-tubular liquid chromatography-electrospray mass spectrometry; ESI, electrospray ionization; oligosaccharide (OS); GlcN, 2-amino-2-deoxy-d-glucose; GlcN3N, β-1’-6 linked 3-diamino-2, 3-dideoxy-D-glucopyranose; PEtn, phosphoethanolamine; Hep, L-glycero-D-manno-heptose; Kdo, 3-deoxy-D-manno-octulosonic residue; Qui*p*3NAcyl, 3-acylamino-3,6-dideoxy-D-glucose; HexNac, hexosamine; deoxyHex, deoxyhexose; Hex, hexose; Qui*p*3NAc, 3-acetamido-3,6-dideoxy-D-glucose; LPS, lipopolysaccharide.

## ACKNOWLEDGEMENTS

The authors wish to thank Ann-Katrin Llarena for her comments and Marja-Liisa Hänninen for providing the strains. This research project was supported by the University of Helsinki research grant n. 313/51/2013. AC was supported by the Microbiology and Biotechnology graduate program (MBDP) from the University of Helsinki. SS and BP were supported by the Biotechnology and Biological Sciences Research Council (BBSRC) grant BB/I02464X/1, and the Medical Research Council (MRC) grants 473 MR/M501608/1 and MR/L015080/1.

## REFERENCES

1. EFSA, and ECDC. 2015. The European Union summary report on trends and sources of zoonoses, zoonotic agents and food-borne outbreaks in 2013. EFSAJournal. 13:3991. doi: 10.2903/j.efsa.2015.3991.

2. Gillespie IA, O’Brien SJ, Frost AF, Adak GK, Horby P, Swan AV, Painter MJ, Neal KR. 2002. A case-case comparison of *Campylobacter coli* and *Campylobacter jejuni* infection: A tool for generating hypotheses. Emerg Infect Dis. 8:937. doi: 10.3201/eid0809.010187.

3. Gürtler M, Alter T, Kasimir S, Fehlhaber K. 2005. The importance of *Campylobacter coli* in human campylobacteriosis: prevalence and genetic characterization. Epidemiol Infect. 133:1081. doi: 10.1017/S0950268805004164.

4. Young, KT, Davis LM, DiRita VJ. 2007. *Campylobacter jejuni:* molecular biology and pathogenesis. Nature Rev Microbiol. 5:665–79.

5. Karlyshev AV, Ketley JM, Wren BW. 2005. The *Campylobacter jejuni* glycome. FEMS Microbiol. Rev. 29:377–390. doi: 10.1016/j.fmrre.2005.01.003.

6. Iwata T, Chiku K, Amano K, Kusumoto M, Ohnishi-Kameyama M, Ono H, Akiba M. 2013. Effects of lipooligosaccharide inner core truncation on bile resistance and chick colonization by *Campylobacter jejuni*. PLoS ONE. 8:e56900.

7. Javed MA, Cawthraw SA, Baig A, Li J, McNally A, Oldfield NJ, Newell DG, Manninga G. 2012. Cj1136 is required for lipooligosaccharide biosynthesis, hyperinvasion, and chick colonization by *Campylobacter jejuni*. Infect Immun. 80:2361–2370.

8. Naito M, Frirdich E, Fields JA, Pryjma M, Li J, Cameron A, Gilbert M, Thompson SA, Gaynor EC. 2010. Effects of sequential *Campylobacter jejuni* 81-176 lipooligosaccharide core truncations on biofilm formation, stress survival, and pathogenesis. J Bacteriol. 192:2182–2192.

9. van Mourik AL, Steeghs L, van Laar J, Meiring HD, Hamstra HJ, van Putten JP, Wösten MM. 2010. Altered linkage of hydroxyacyl chains in lipid A of *Campylobacter jejuni* reduces TLR4 activation and antimicrobial resistance. J Biol Chem. 285:15828–15836. doi: 10.1074/jbc.M110.102061.

10. Stephenson HN, John CM, Naz N, Gundogdu O, Dorrell N, Wren BW, Jarvis GA, Bajaj-Elliott M. 2013. *Campylobacter jejuni* lipooligosaccharide sialylation, phosphorylation, and amide/ester linkage modifications fine-tune human toll-like receptor 4 activation. J Biol Chem 288:19661–19672. doi: 10.1074/jbc.M113.468298.

11. Kuijf ML, Samsom JN, van Rijs W, Bax M, Huizinga R, Heikema AP, van Doorn PA, van Belkum A, van Kooyk Y, Burgers PC, Luider TM, Endtz HP, Nieuwenhuis EE, Jacobs BC. 2010. TLR4-mediated sensing of *Campylobacter jejuni* by dendritic cells is determined by sialylation. J Immunol 185:748–755. doi: 10.4049/jimmunol.0903014.

12. Bax M, Kuijf ML, Heikema AP, van Rijs W, Bruijns SC, García-Vallejo JJ, Crocker PR, Jacobs BC, van Vliet SJ, van Kooyk Y. 2011. *Campylobacter jejuni* lipooligosaccharides modulate dendritic cell-mediated T cell polarization in a sialic acid linkage-dependent manner. Infect Immun. 79:2681–2689. doi: 10.1128/IAI.00009-11.

13. Huizinga R, van Rijs W, Bajramovic JJ, Kuijf ML, Laman JD, Samsom JN, Jacobs BC. 2013. Sialylation of *Campylobacter jejuni* endotoxin promotes dendritic cell-mediated B cell responses through CD14-dependent production of IFN-ß and TNF-a. J Immunol. 191:5636–5645. doi: 10.4049/jimmunol.1301536.

14. Huizinga R, Easton AS, Donachie AM, Guthrie J, van Rijs W, Heikema A, Boon L, Samsom JN, Jacobs BC, Willison HJ, Goodyear CS. 2012. Sialylation of *Campylobacter jejuni* lipo-oligosaccharides: Impact on phagocytosis and cytokine production in mice. PLoS ONE. 7:e34416.

15. Yuki N, Susuki K, Koga M, Nishimoto Y, Odaka M, Hirata K, Taguchi K, Miyatake T, Furukawa K, Kobata T, Yamada M. 2004. Carbohydrate mimicry between human ganglioside GM1 and *Campylobacter jejuni* lipooligosaccharide causes Guillain-Barré syndrome. Proc Natl Acad Sci USA. 101:11404–11409. doi: 10.1073/pnas.0402391101.

16. Gilbert M, Karwaski M, Bernatchez S, Young NM, Taboada E, Michniewicz J, Cunningham A, Wakarchuk WW. 2002. The genetic bases for the variation in the lipo-oligosaccharide of the mucosal pathogen, *Campylobacter jejuni*. J Biol Chem. 277:327–337. doi: 10.1074/jbc.M108452200.

17. Dzieciatkowska M, Brochu D, van Belkum A, Heikema AP, Yuki N, Houliston RS, Richards JC, Gilbert M, Li J. 2007. Mass spectrometric analysis of intact lipooligosaccharide: Direct evidence for O-acetylated sialic acids and discovery of O-linked glycine expressed by *Campylobacter jejuni*. Biochemistry. 46:14704–14714.

18. Chiu CP, Watts AG, Lairson LL, Gilbert M, Lim D, Wakarchuk WW, Withers SG, Strynadka NC. 2004. Structural analysis of the sialyltransferase CstII from *Campylobacter jejuni* in complex with a substrate analog. Nat Struct Mol Biol. 11:163–170.

19. Parker CT, Gilbert M, Yuki N, Endtz HP, Mandrell RE. 2008. Characterization of lipooligosaccharide-biosynthetic loci of *Campylobacter jejuni* reveals new lipooligosaccharide classes: evidence of mosaic organizations, J Bacteriol 190:5681–5689.

20. Linton D, Gilbert M, Hitchen PG, Dell A, Morris HR, Wakarchuk WW, Gregson NA, Wren BW. 2000. Phase variation of a β-1,3 galactosyltransferase involved in generation of the ganglioside GM1-like lipo-oligosaccharide of *Campylobacter jejuni*. Mol Microbiol. 37:501–514. doi: 10.1046/j.1365-2958.2000.02020.x.

21. Revez, J, Hänninen M. 2012. Lipooligosaccharide locus classes are associated with certain *Campylobacter jejuni* multilocus sequence types. Eur J Clin Microbiol Infect Dis. 31:2203–2209.

22. Skarp-de Haan CP, Culebro A, Schott T, Revez J, Schweda E, Hänninen M, Rossi M. 2014. Comparative genomics of unintrogressed *Campylobacter coli* clades 2 and 3. BMC Genomics. 15:129.

23. Richards VP, Lefébure T, Pavinski Bitar PD, Stanhope MJ. 2013. Comparative characterization of the virulence gene clusters (lipooligosaccharide [LOS] and capsular polysaccharide [CPS]) for Campylobacter coli, Campylobacter jejuni subsp. jejuni and related Campylobacter species. Infect Genet Evol. 14:200–213. doi: 10.1016/j.meegid.2012.12.010.

24. Aspinall GO, McDonald AG, Pang H, Kurjanczyk LA, Penner JL. 1993. Lipopolysaccharide of *Campylobacter coli* serotype O:30. Fractionation and structure of liberated core oligosaccharide. J Biol Chem. 268:6263–6268.

25. Aspinall GO, McDonald AG, Pang H, Kurjanczyk LA, Penner JL. 1993. An antigenic polysaccharide from *Campylobacter coli* serotype O:30. Structure of a teichoic acid-like antigenic polysaccharide associated with the lipopolysaccharide. J Biol Chem. 268:18321–18329.

26. Beer W, Adam M, Seltmann G. 1986. Monosaccharide composition of lipopolysaccharides from *Campylobacter jejuni* and *Campylobacter coli*. J Basic Microbiol. 26:201–204. doi: 10.1002/jobm.3620260405.

27. Naess V, Hofstad T. 1984. Chemical studies of partially hydrolysed lipopolysaccharides from four strains of *Campylobacter jejuni* and two strains of *Campylobacter coli*. J Gen Microbiol. 130:2783–2789.

28. Sheppard SK, Didelot X, Jolley KA, Darling AE, Pascoe B, Meric G, Kelly DJ, Cody A, Colles FM, Strachan NJ, Ogden ID, Forbes K, French NP, Carter P, Miller WG, McCarthy ND, Owen R, Litrup E, Egholm M, Affourtit JP, Bentley SD, Parkhill J, Maiden MC, Falush D. 2013. Progressive genome-wide introgression in agricultural *Campylobacter coli*. Mol Ecol. 22:1051–1064. doi: 10.1111/mec.12162.

29. Sheppard SK, Maiden MC. 2015. The evolution of *Campylobacter jejuni* and *Campylobacter coli*. Cold Spring Harb Perspect Biol. 7: doi: 10.1101/cshperspect.a018119.

30. Olkkola SH, Juntunen P, Heiska H, Hyytiäinen H, Hänninen M. 2010. Mutations in the *rpsL* gene are involved in streptomycin resistance in *Campylobacter coli*. Microb Drug Resist. 16:105. doi: 1089/mdr.2009.0128.

31. Juntunen P, Olkkola S, Hänninen M. 2011. Longitudinal on-farm study of the development of antimicrobial resistance in *Campylobacter coli* from pigs before and after danofloxacin and tylosin treatments. Vet Microbiol. 150:322–330.

32. Juntunen P, Heiska H, Olkkola S, Myllyniemi AL, Hänninen ML. 2010. Antimicrobial resistance in *Campylobacter coli* selected by tylosin treatment at a pig farm. Vet Microbiol. 146:90–97.

33. Lehtopolku M, Kotilainen P, Haanperä-Heikkinen M, Nakari U, Hänninen M, Huovinen P, Siitonen A, Eerola E, Jalava J, Hakanen AJ. 2011. Ribosomal mutations as the main cause of macrolide resistance in *Campylobacter jejuni* and *Campylobacter coli*. Antimicrob Agents Chemother. 55:5939–5941. doi: 10.1128/AAC.00314-11.

34. Kärenlampi R, Rautelin H, Schönberg-Norio D, Paulin L, Hänninen M. 2007. Longitudinal study of Finnish *Campylobacter jejuni and C. coli* isolates from humans, using multilocus sequence typing, including comparison with epidemiological data and isolates from poultry and cattle. Appl Environ Microbiol. 73:148–155. doi: 10.1128/AEM.01488-06.

35. Llarena AK, Skarp-de Haan CP, Rossi M, Hänninen M. 2015. Characterization of the *Campylobacter jejuni* population in the barnacle geese reservoir. Zoonoses and Public Health. 62:209–221. doi: 10.1111/zph.12141.

36. Hänninen M, Pajarre S, Klossner M, Rautelin H. 1998. Typing of human *Campylobacter jejuni* isolates in Finland by pulsed-field gel electrophoresis. J Clin Microbiol. 36:1787–1789.

37. Darling AE, Mau B, Perna NT. 2010. progressiveMauve: multiple genome alignment with gene gain, loss and rearrangement. PLoS One. 5:e11147.

38. Bankevich A, Nurk S, Antipov D, Gurevich AA, Dvorkin A, Kulikov AS, Lesin VM, Nikolenko SI, Pham S, Prjibelski AD, Pyshkin AV, Sirotkin AV, Vyahhi N, Tesler G, Alekseyev MA, Pevzner PA. 2012. SPAdes: A new genome assembly algorithm and its applications to single-cell sequencing. J Comput Biol. 19:455–477. doi: 10.1089/cmb.2012.0021.

39. Aziz RK, Bartels D, Best AA, DeJongh M, Disz T, Edwards RA, Formsma K, Gerdes S, Glass EM, Kubal M, Meyer F, Olsen GJ, Olson R, Osterman AL, Overbeek RA, McNeil LK, Paarmann D, Paczian T, Parrello B, Pusch GD, Reich C, Stevens R, Vassieva O, Vonstein V, Wilke A, Zagnitko O. 2008. The RAST Server: Rapid Annotations using Subsystems Technology. BMC Genomics. 9:1–15. doi: 10.1186/1471-2164-9-75.

40. Rutherford K, Parkhill J, Crook J, Horsnell T, Rice P, Rajandream M, Barrell B. 2000. Artemis: sequence visualization and annotation. Bioinformatics. 16:944–945. doi: 10.1093/bioinformatics/16.10.944.

41. Carver T, Berriman M, Tivey A, Patel C, Böhme U, Barrell BG, Parkhill J, Rajandream MA. 2008. Artemis and ACT: viewing, annotating and comparing sequences stored in a relational database. Bioinformatics. 24:2672–2676. doi: 10.1093/bioinformatics/btn529.

42. Altschul SF, Madden TL, Schäffer AA, Zhang J, Zhang Z, Miller W, Lipman DJ. 1997. Gapped BLAST and PSI-BLAST: a new generation of protein database search programs. Nucleic Acids Research. 25:3389–3402. doi: 10.1093/nar/25.17.3389.

43. Enright AJ, van Dongen S, Ouzounis CA. 2002. An efficient algorithm for large-scale detection of protein families. Nucleic Acids Res. 30:1575–1584. doi: 10.1093/nar/30.7.1575.

44. Ekseth OK, Kuiper M, Mironov V. 2013. orthAgogue: an agile tool for the rapid prediction of orthology relations. Bioinformatics. 30: 734–736. doi: 10.1093/bioinformatics/btt582.

45. Edgar RC, Sjolander K. 2004. A comparison of scoring functions for protein sequence profile alignment. Bioinformatics. 20:1301–1308

46. Edgar RC. 2004. MUSCLE: a multiple sequence alignment method with reduced time and space complexity. BMC Bioinformatics. 5:1–19. doi: 10.1186/1471-2105-5-113.

47. Abascal F, Zardoya R, Telford MJ. 2010. TranslatorX: multiple alignment of nucleotide sequences guided by amino acid translations. Nucleic Acids Res. 8:W7–W13. doi: 10.1093/nar/gkq291.

48. Tamura K, Stecher G, Peterson D, Filipski A, Kumar S. 2013. MEGA6: Molecular evolutionary genetics analysis version 6.0. Mol Biol Evol. 30: 2725–2729. doi: 10.1093/molbev/mst197.

49. Revez J, Rossi M, Ellström P, de Haan CP, Rautelin H, Hänninen M. 2011. Finnish *Campylobacter jejuni s*trains of multilocus sequence type ST-22 complex have two lineages with different characteristics. PLoS ONE. 6:e26880.

50. Dzieciatkowska M, Liu X, Heikema AP, Houliston RS, van Belkum A, Schweda EK, Gilbert M, Richards JC, Li J. 2008. Rapid method for sensitive screening of oligosaccharide epitopes in the lipooligosaccharide from *Campylobacter jejuni* strains isolated from Guillain-Barré syndrome and Miller Fisher syndrome patients. J Clin Microbiol. 46:3429–3436. doi: 10.1128/JCM.00681-08.

51. Moran AP, Zähringer U, Seydel U, D. Scholz, Stütz P, Rietschel ET. 1991. Structural analysis of the lipid A component of *Campylobacter jejuni* CCUG 10936 (serotype O:2) lipopolysaccharide. Eur J Biochem. 198:459–469. doi: 10.1111/j.1432-1033.1991.tb16036.x.

52. Moran AP, Rietschel ET, Kosunen TU, Zähringer U. 1991. Chemical characterization of *Campylobacter jejuni* lipopolysaccharides containing N-acetylneuraminic acid and 2,3-diamino-2,3-dideoxy-D-glucose. J Bacteriol. 173:618–626.

53. Moran AP. 1997. Structure and conserved characteristics of *Campylobacter jejuni* lipopolysaccharides. J Infect Dis. 176:S115–S121. doi: 10.1086/513781.

54. Stahl M, Ries J, Vermeulen J, Yang H, Sham HP, Crowley SM, Badayeva Y, Turvey SE, Gaynor EC, Li X. 2014. A novel mouse model of *Campylobacter jejuni* gastroenteritis reveals key pro-inflammatory and tissue protective roles for toll-like receptor signaling during Infection. PLoS Pathog. 10:e1004264. doi: doi: 10.1371/journal.ppat.1004264.

55. Bereswill S, Fischer A, Plickert R, Haag L, Otto B, Kühl AA, Dashti JI, Zautner AE, Muñoz M, Loddenkemper C, Groß U, Göbel UB, Heimesaat MM. 2011. Novel murine infection models provide deep insights into the "ménage à trois" of *Campylobacter jejuni*, microbiota and host innate immunity. PLoS ONE. 6:e20953.

56. Roux F, Sproston E, Rotariu O, MacRae M, Sheppard SK, Bessell P, Smith-Palmer A, Cowden J, Maiden MC, Forbes K, Strachan NJ. 2013. Elucidating the aetiology of human *Campylobacter coli* infections. PLoS ONE. 8:e64504.

57. Sweet CR, Ribeiro AA, Raetz CR. 2004. Oxidation and transamination of the 3?-Position of UDP-N-acetylglucosamine by enzymes from *Acidithiobacillus ferrooxidans*. J Biol Chem. 279:25400–25410. doi: 10.1074/jbc.M400596200.

58. Jansonius JN. 1998. Structure, evolution and action of vitamin B6-dependent enzymes. Curr Opin Struct Biol. 8:759–769.

59. Altman E, Schäffer C, Brisson JR, Messner P. 1995. Characterization of the glycan structure of a major glycopeptide from the surface layer glycoprotein of *Clostridium thermosaccharolyticum* E207-71. Eur. J. Biochem. 229:308–315. doi: 10.1111/j.1432-1033.1995.0308l.x.

60. Ovchinnikova OG, Rozalski A, Liu B, Knirel YA. 2013. O-antigens of bacteria of the genus *Providencia*: Structure, serology, genetics, and biosynthesis. Biochemistry (Mosc). 78:798–817. doi: 10.1134/S0006297913070110.

61. Veremeichenko S, Zdorovenko GM. 2004. Structure and properties of the lipopolysaccharide of *Pseudomonas fluorescens* IMV 2366 (biovar III)]. Microbiology. 73:312–319.

62. Holden KM, Gilbert M, Coloe PJ, Li J, Fry BN. 2012. The role of WlaRG, WlaTB and WlaTC in lipooligosaccharide synthesis by *Campylobacter jejuni* strain 81116. Microb Pathog. 52:344–352. doi: 10.1016/j.micpath.2012.03.004.

63. Godschalk PC, Gilbert M, Jacobs BC, Kramers T, Tio-Gillen AP, Ang CW, Van den Braak N, Li J, Verbrugh HA, van Belkum A, Endtz HP. 2006. Co-infection with two different *Campylobacter jejuni* strains in a patient with the Guillain-Barré syndrome. Microb Infect. 8:248–253.

64. Aspinall GO, Lynch CM, Pang H, Shaver RT, Moran AP. 1995. Chemical structures of the core region of *Campylobacter jejuni* O:3 lipopolysaccharide and an associated polysaccharide. Eur J Biochem. 231:570–578. doi: 10.1111/j.1432-1033.1995.tb20734.x.

65. Seltmann G, Beer W. 1985. Vorkommen von 3-Amino-3,6-didesoxy-D-glucose in einem Lipopolysaccharid von *Campylobacter coli*. J Basic Microbiol. 25:551–552.

66. Pföstl A, Zayni S, Hofinger A, Kosma P, Schäffer C, Messner P. 2008. Biosynthesis of dTDP-3-acetamido-3,6-dideoxy-?-D-glucose. Biochem J. 410:187–194. doi: 10.1042/BJ20071044.

67. Bernatchez S, Szymanski CM, Ishiyama N, Li J, Jarrell HC, Lau PC, Berghuis AM, Young NM, Wakarchuk WW. 2005. A single bifunctional UDP-GlcNAc/Glc 4-epimerase supports the synthesis of three cell surface glycoconjugates in *Campylobacter jejuni*. J Biol Chem. 280:4792–4802. doi: 10.1074/jbc.M407767200.

68. Ma B, Simala-Grant JL, Taylor DE. 2006. Fucosylation in prokaryotes and eukaryotes. Glycobiology. 16:158R–184R. doi: 10.1093/glycob/cwl040.

69. Knirel, Y. 2011. Knirel, Y., 2011. Structure of O-antigens. in: Knirel, Y.A., Valvano, M.A. (Eds.), Bacterial Lipopolysaccharides, Springer, Vienna, Austria, 41-115., p. 41–115. In Y. A. Knirel and M. A. Valvano (eds.), Bacterial Lipopolysaccharides. Springer, Vienna, Austria.

70. Hanniffy OM, Shashkov AS, Moran AP, Prendergast MM, Senchenkova SN, Knirel YA, Savage AV. 1999. Chemical structure of a polysaccharide from Campylobacter jejuni 176.83 (serotype O:41) containing only furanose sugars. Carbohydr Res. 319:124–132.

71. Moran AP, O’Malley DT, Kosunen TU, Helander IM. 1994. Biochemical characterization of *Campylobacter fetus* lipopolysaccharides. Infect Immun. 62:3922–3929.

